# Rapid and sensitive quantitation of glucose and glucose phosphates derived from storage carbohydrates using gas chromatography mass spectrometry

**DOI:** 10.1101/645556

**Authors:** Lyndsay E.A. Young, Corey O. Brizzee, Jessica K. A. Macedo, Matthew S. Gentry, Ramon C. Sun

## Abstract

Glycogen is the primary storage carbohydrate in mammals and it is synthesized in most tissues. Glycogen contains covalently attached phosphate groups on hydroxyls of glucose units. The addition of phosphate modulates branching pattern, granular size, and crystallinity of a glycogen molecule, which all impact its accessibility to glycogen interacting enzymes during catabolism. As glycogen architecture modulates its role in metabolism, it is essential to accurately evaluate and quantify phosphate content in glycogen. Simultaneous quantitation of glucose and its phosphate esters is challenging and requires an assay with high sensitivity and a robust dynamic range. Currently, this method is lacking in the field. Herein, we describe a highly-sensitive method for the detection of both glycogen-derived glucose and glucose-phosphate esters utilizing gas-chromatography coupled mass spectrometry. Using this method, we observed higher glycogen levels in the liver compared to skeletal muscle, but skeletal muscle contained much more phosphate esters. These results confirm previous findings and establish the validity of the method. Importantly, this method can detect femtomole levels of glucose and glucose phosphate esters within an extremely robust dynamic range with excellent accuracy and reproducibility. The method can also be easily adapted for the quantification of glucose from plant starch, amylopectin or other biopolymers as well as covalently attached phosphate within them.

## Introduction

Glycogen is a branched polymer of glucose moieties that functions as an energy reserve in mammals. Glycogen is found in most tissues, including liver (1,2), muscle (3), kidney (4), brain (5), and white blood cells (6). The synthesis and breakdown of glycogen involves several enzymes and regulatory proteins (7). Glycogen is comprised of glucose moieties joined by α-1,4-glycosidic linkages, formed by glycogen synthase, and branches occurring every 12-14 units via α-1,6-glycosidic branches, formed by glycogen branching enzyme. This unique organization allows cells to store up to ≈55,000 glucose units in a water-soluble form with high packing density for maximum storage (8). During glycogenolysis (glycogen breakdown), glycogen phosphorylase (GP) (9) releases glucose-1-phosphate molecules to fuel a wide range of metabolic processes (10). Glycogen synthesis and degradation either consumes or produces free glucose-6-phosphate (G6P), a key metabolite essential for energy production, lipid generation, and nucleotide biosynthesis important for cellular physiology.

Glycogen contains phosphate monoester groups covalently attached to the C2-, C3-, and C6-position of glucose hydroxyls (11–13). Glycogen-bound phosphate modulates glycogen architecture through branching and chain length that define glycogen granular size, crystallinity (solubility), and its accessibility to glycogen interacting enzymes (14–17). Liver glycogen has an average chain length of 13 residues (11,18) and approximately 1 phosphate per 1,000-10,000 glucose residues (11,12). These properties allow liver glycogen to maintain maximum solubility for rapid turn-over during periods of starvation. Muscle glycogen is architecturally distinct from liver glycogen. Muscle glycogen exhibits increases in phosphate esters, altered branching pattern compared to liver glycogen, and it is a fuel storage for the fight-or-flight response and does not contribute to the regulation of blood glucose (12,19). While the mechanism of phosphate incorporation into glycogen remains unresolved, glycogen phosphorylation is intimately linked with the biophysical properties and biological utilization of glycogen (11,20–22). The importance of regulating glycogen architecture and phosphorylation is highlighted in Lafora disease (LD). LD is an early onset neurodegenerative disease resulting from the accumulation of hyperphosphorylated, aberrant glycogen aggregates that drive disease pathogenesis in the form of myoclonic seizures, neuroinflammation, and premature death (23–25).

Glucose-3-phosphate (G3P) and glucose-6-phosphate (G6P) are the hydrolyzed monomeric forms that represent glycogen phosphate content. Although G3P and G6P are biochemically similar, G6P is a naturally occurring free metabolite that participates in glycolysis and the pentose phosphate pathway. G6P can be measured via spectrophotometric methods utilizing G6P dehydrogenase and NADP/NADPH conversion (26). Conversely, an enzyme that utilizes G3P has not been discovered. Therefore, the two cannot be distinguished by conventional biochemical assays. Currently, the technologies that can unambiguously distinguish G3P and G6P are 2D-nuclear magnetic resonance (NMR) and capillary electrophoresis (CE). 2D-NMR is time consuming, requires a minimum of micrograms to milligrams of glycogen for accurate assignment, and it is challenging to adapt for routine laboratory analysis (27). Currently, fluorescence-assisted capillary electrophoresis (FACE) is the gold standard to quantify C3 and C6 phosphorylation of glucose residues in starch (28). Starch is hydrolyzed and the glucose moieties are conjugated to 8-aminopyrene-1,3,6-trisulfonic acid (APTS). The APTS-conjugated glucose moieties can then be quantified using CE with accurate measurements as low as 1 glucose phosphate/100-1,000 glucose residues in plant starch. However, mammalian glycogen often contains much less glucose-bound phosphate, on the order of 10-to 100-fold less, which is below the dynamic range for CE. In addition, the presence of free APTS and inherent variation in retention time both add additional difficulties for batch processing, especially with low phosphate levels. Therefore, a higher-throughout, more accessible assay is needed that yields a more robust dynamic range with higher reproducibility and simultaneous quantitation of glucose, G3P, and G6P.

Herein, we introduce a workflow for the extraction and quantitation of glycogen and its phosphate esters. Glycogen was hydrolyzed to glucose, G3P, and G6P monomers and separated by gas-chromatography coupled to a highly sensitive mass spectrometry (GCMS) that can analyze a robust dynamic range and sensitivity for all three metabolites. With the addition of an auto-sampler, this GCMS-based method can profile up to 120 samples/day with accuracy, reliability, and reproducibility for routine interrogation of glucose moieties from mammalian glycogen and other glucose-based biopolymers.

## Results

### Chemical derivatization of Glucose, G3P, and G6P

Previous work utilizing multiple tissues demonstrated the reliable and sensitive separation of free sugar-phosphates after chemical derivatization for metabolomics applications (29). We hypothesized that this method could also separate G3P and G6P following hydrolysis of glycogen. First, we performed a two-step derivatization procedure using analytical standards to convert them to volatile trimethylsilyl (TMS) derivatives that can be detected by a mass spectrometer (30,31). In the first step, the methoxylamination reaction replaces the oxygen atom of the alpha-carbonyl groups with methoxyamine (MEOX). Then silylation is performed in the second derivatization step using N-methyl-N-trimethylsilylation (MSTFA) to introduce trimethylsilyl groups to the remaining 6 carboxyl groups, replacing the acidic hydrogens (**Fig. 1A-C**). Methoxyamination generates both the syn- and ant-forms of MEOX at the alpha-carbonyl group, therefore resulting in the formation of a second much smaller peak with a small retention time shift that does not affect the quantification step (**Fig. 2B**). Trimethylsilylation of metabolites are subjected to electron ionization (EI), where electrons fragment the molecules so that they can be registered by the mass spectrometer detector as molecular ions. The fragmentation pattern is consistent and reproducible across different GCMS platforms (**Fig. 1A-C**). In selected ion monitoring (SIM) mode, only specific fragment ions are detected by the mass spectrometer, significantly improving accuracy and sensitivity. Fragment ions (*m/z*) 319, 364 (glucose), 315, 357 and 387 (G3P and G6P) were the major ions produced from EI-MS (**Fig. 1A-C**), and were used for the rest of the study in SIM mode to improve sensitivity of the GCMS.

**Fig. 1.**
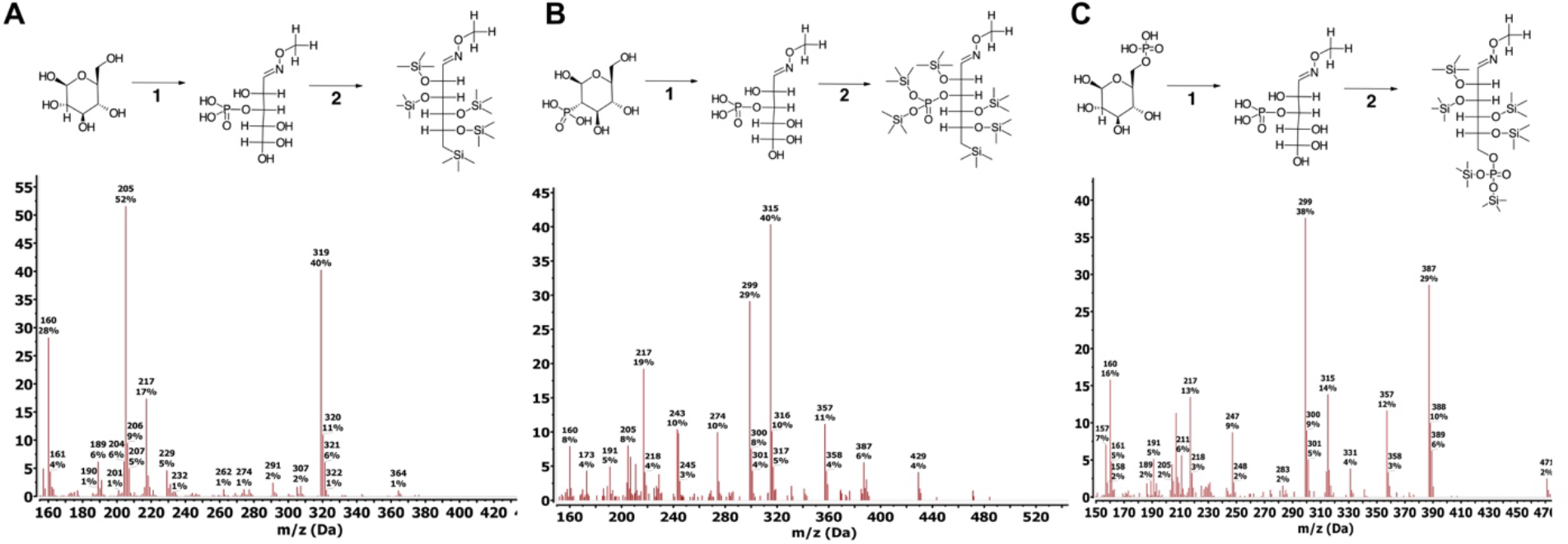
Derivatization and fragmentation pattern of glucose, G3P, G6P. Trimethylsilylation of analytical grade standards of glucose (**A**), G3P (**B**), and G6P (**C**) (−6TMS;1MEOX). Two μmoles of each standard were derivatized with 20mg/ml MEOX in pyridine for 1-hour (reaction 1), followed by silylation by MSTFA for 1-hour (reaction 2). Both reaction steps took place in a 60° C dry heat block. Fragmentation pattern for each silylated standard was obtained on a single quadrupole mass spectrometer with an electron ionization (EI) energy of 70 eV, mass range of 30-650 AMU, and 1.47 scan/s.

### Gas chromatography separation of Glucose, G3P, and G6P

G3P and G6P share similar chemical properties and specific detection has proven to be difficult. We hypothesized that the unique TMS derivatized form of G3P and G6P could be separated via gas chromatography (GC) given the appropriate temperature gradient. Each TMS derivative was analyzed separately using an adapted-Fiehn metabolomics GC method to confirm retention time (**Fig. 2A**) (32,33). Glucose-, G3P-, and G6P-6TMS;1MEOX eluted at 17.6, 21.2 and 21.7 minutes respectively (± 0.05sec) and confirmed the utility of the GC to separate glucose and glucose phosphate esters (**Fig. 2B**). We identified the optimal separation temperature as 180-280° C with a ramp speed of 1” C/minute. These parameters were adapted and reduced the processing time to 12 minutes to provide a more rapid separation of the glucose moieties (**Fig. 2C**). We tested the ability of the rapid method to separate all three TMS derivatives by combining 1 nmol of each standard into one sample and analyzed the combined standards by the GCMS method. The retention times for glucose, G3P, and G6P were 6.25, 7.23, and 7.46 minutes (± 0.05sec), respectively (**Fig. 2D**). Thus, the rapid method yields a clear separation between glucose and each of the two phosphate esters.

**Fig. 2.**
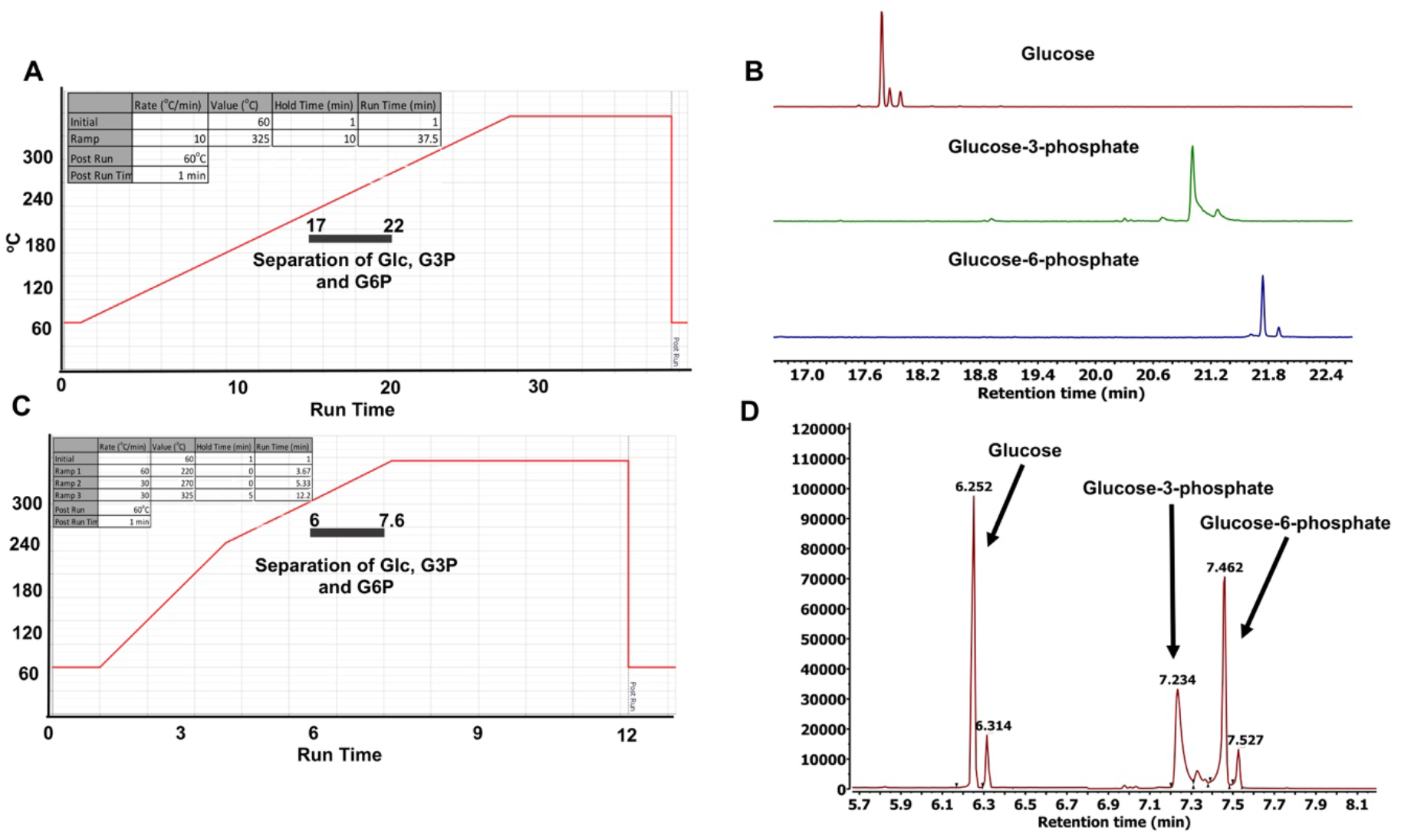
Standard and rapid gas chromatography separation of glucose, G3P, G6P. (**A**) Temperature gradient for standard separation of glucose, G3P, and G6P: Initial temperature was 60° C, held for 1 minute, rising at 10° C/minute to 325° C, held for 10 minutes. Total run time: 37.5 minutes. Grey bar indicates window of separation. (**B**) Stacked chromatography spectra for silylated glucose, G3P, and G6P using the temperature setting in (**A**). Twenty nmoles of each silylated standard were injected into the GC column. (**C**) Temperature gradient setting for rapid separation of glucose, G3P, and G6P: initial temperature was 60° C, held for 1 minute, rising at 60° C/minute to 220° C, continued rising at 30° C/minute to 270° C, and finished rising at 30° C/minute to 325° C, held for 5 minutes. Total run time: 12.2 minutes. Grey bar indicates window of separation. (**D**) Chromatography separation for silylated glucose, G3P, and G6P combined into one sample using the temperature setting in (**C**). Two nmoles of each silylated standard were injected into the GC column.

### Dynamic range of Glucose, G3P, and G6P

Mammalian glycogen contains 1 phosphate ester/1000-10,000 glucose residues, approximately 10-to 100-fold lower glucose-bound phosphate than potato starch (11,12). To simultaneously detect glucose, G3P, and G6P monomers from glycogen, an assay with a robust and dynamic range is needed. Following the successful separation of TMS derivatized glucose moieties, we proceeded to determine the sensitivity and dynamic range for each molecule.

We tested the limit of detection and dynamic range using both splitless and the 10:1 split mode of the GC inlet. In split mode, the limit of detection was 10 pmol to 10 μmol for glucose, G3P, and G6P (**Fig. 3A-C**). In splitless mode, the limit of detection was 10 fmol for all three compounds with a dynamic range of 10 fmol to 10 pmol (**Fig. 3D-F**). We did not test the range any lower as it is beyond the physiological range and potentially damaging to the ion source of the mass spectrometer. Cumulatively, these data confirm that GCMS analysis is extremely versatile for the separation of glucose, G3P, and G6P and it possesses a robust dynamic range that is well-suited for the analysis of a wide range of biopolymers.

**Fig. 3.**
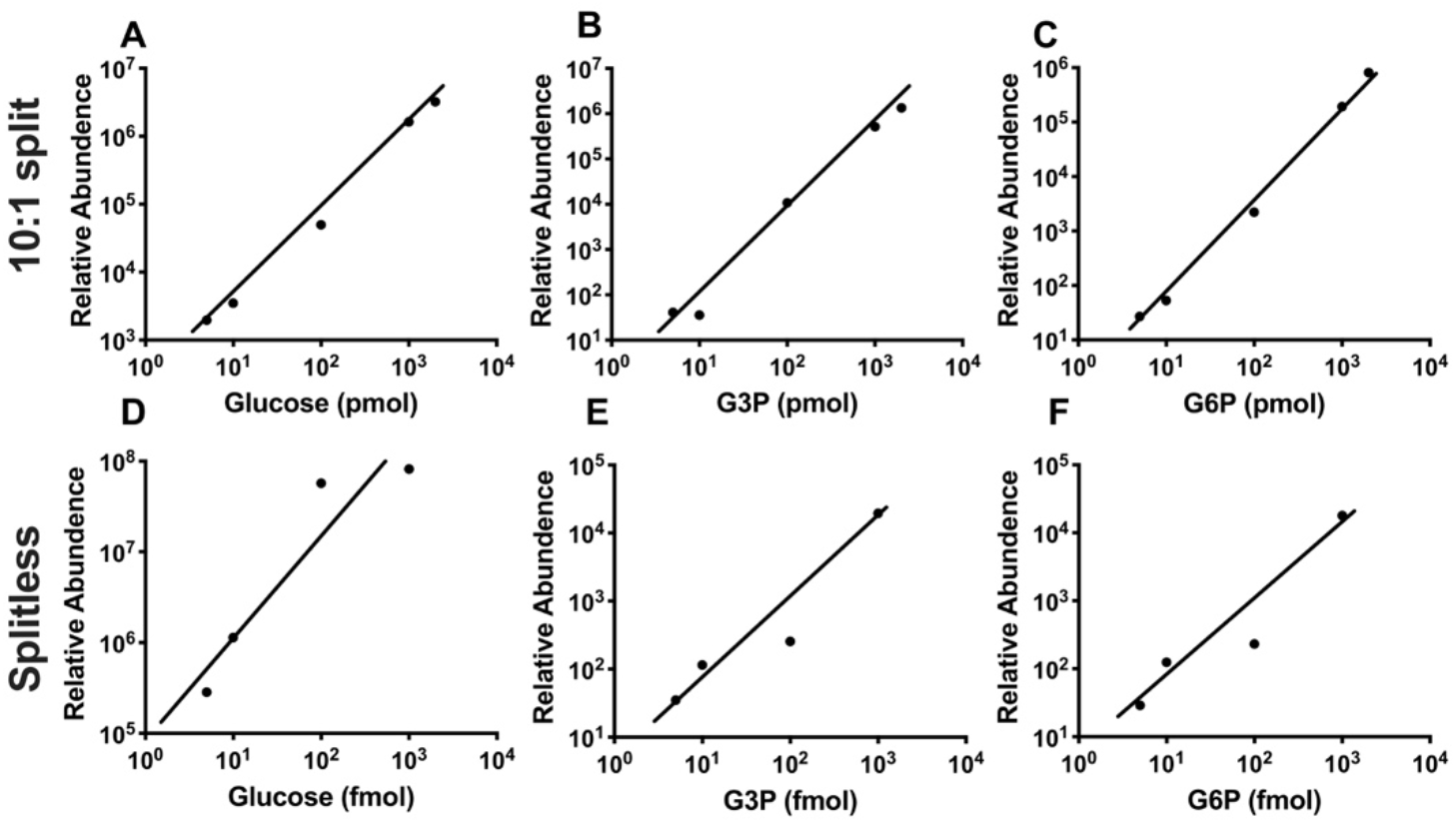
Dynamic range of GCMS analysis of glucose, G3P, and G6P. Silylated glucose (**A**), G3P (**B**), and G6P (**C**) standards at multiple concentrations were injected into the GC using split mode with a ratio of 10:1. For boosted sensitivity to detect samples with low glycogen content, lower concentrations of silylated glucose (**D**), G3P (**E**), and G6P (**F**) standards were analyzed on GCMS using the splitless injection mode.

### Quantitation of glycogen and phosphate from mouse skeletal muscle and liver

Glycogen architecture gives rise to unique physiochemical properties that cause barriers to purifying it. The current method of glycogen purification from tissue uses 10% trichloroacetic acid (TCA) for low phosphate glycogen such as liver, and boiling the tissue in potassium hydroxide for muscle or other high phosphate containing glycogen molecules. Both techniques require 10g-100g of tissue material due to low yield, and they introduce variations in extraction efficiency and yield ambiguity in purity. In this study, we adapted a 2-step purification procedure that is highly efficient, and yields glycogen from as low as 20mg input from multiple tissues.

First, mouse liver and skeletal muscle were pulverized to 10 μm particles (single cell volume) using a liquid nitrogen magnetic assisted tissue-grinding mill (**Fig. 4A**). After pulverization, 20mg of tissue was used for the glycogen purification method. Free polar metabolites and lipids were removed by the addition of polar and organic solvents, 50% methanol:chloroform (1:1). The insoluble glycogen-containing layer was collected and washed with 50% methanol and allowed to dry. The glycogen was extracted with the addition of 10% trichloroacetic acid followed by vigorous mixing. The glycogen-containing TCA fraction was then separated from other insoluble material by centrifugation, and the glycogen was dried by vacuum centrifuge at 10^−3^ mBar, hydrolyzed to monomers by first resuspending in diH20 followed by the addition of an equal part of 2N HCl and the reaction was carried out at 95°C for 2 hours. Hydrolysis was quenched with 100% methanol, 40uM L-norvaline was added (as an internal control), and the sample was incubated on ice for 30 minutes. The supernatant was collected following a 10-minute 15,000 rpm spin at 4°C and dried by vacuum centrifuge at 10^−3^ mBar. The dried glucose moieties were derivatized by MEOX and MSTFA as defined above, and then analyzed by GCMS (**Fig. 4B**). The polar fraction and the last wash were also analyzed as positive and negative controls (blank), respectively. The analysis found that liver glycogen contains 18±3 μmol of glucose, 4±1 nmol of G3P, and 5±1 nmol of G6P per g of wet tissue weight (**Fig. 4C**). Alternatively, muscle stores lower levels of glycogen (1±0.12 μmol glucose/g of wet tissue weight), yet muscle glycogen contains higher levels of phosphate esters with 12±1 nmol G3P and G6P per g of wet tissue weight (**Fig. 4D and E**). These levels are in the ranges of previously published results and confirm previous findings that demonstrated higher phosphate content in muscle glycogen (11,12,34–37). Finally, we observed G6P in both polar fractions of muscle and liver extracts, but G3P was not detected in the fractions as expected.

**Fig. 4.**
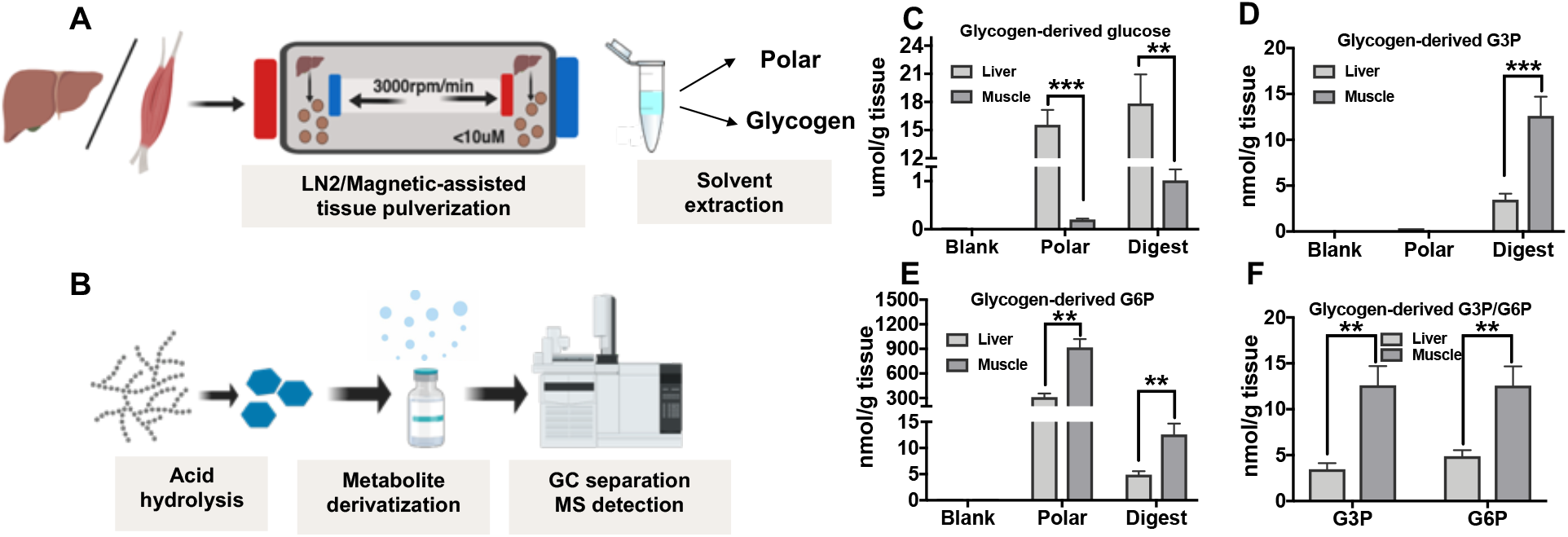
Extraction and GCMS analysis of liver and muscle glycogen. (**A**) Schematics of glycogen extraction: mouse liver or skeletal muscle were milled to 10μm particles by liquid N_2_ Freezer/Mill Cyrogenic Grinder magnetic assisted tissue-grinding mill, followed by polar and organic solvent removal of free polar metabolites and lipids. Glycogen was then extracted by 10% TCA. (**B**) Schematics of glycogen preparation for GCMS analysis: isolated glycogen was hydrolyzed to monomers by mild hydrolysis, derivatized by MEOX and MSTFA, and analyzed by GCMS. (**C**) Quantitation of liver or muscle derived glucose (**C**), G3P (**D**), and G6P (**E**), the third wash of tissue pellet is served as blank, and free polar metabolite fraction serves as negative control for G3P and a positive control for G6P. (**F**) Muscle contains higher G3P and G6P than liver when standardized to tissue weight. * *p* ≤ 0.05, ** *p* ≤ 0.01, *** *p* ≤ 0.001; two-tailed *t*-test.

While the mechanism of glycogen phosphorylation is unresolved, reversible starch phosphorylation is well-defined and is integral for starch metabolism (38–40). Transitory starch is synthesized during the day and degraded at night, and reversible starch phosphorylation is necessary for efficient degradation. Glucan water dikinase (GWD) phosphorylates hydroxyls at the C6-position of glucose moieties on the outer surface of starch (41) (42). This phosphorylation event triggers phosphorylation at the C3-position by phosphoglucan water dikinase (PWD) (43,44). These coordinated phosphorylation events allow amylases to more efficiently liberate glucose from starch until they reach the phosphate moieties. The glucan phosphatases starch excess 4 (SEX4) and like-sex four2 (LSF2) then liberate the phosphate so that the process continues (45,46). Plants lacking GWD activity lack starch phosphorylation at the C6- and C3-position (44).

To test the versatility of this method, we performed a similar GCMS analysis on starch and glycogen. We analyzed wild-type (WT) starch from potato, starch from *Arabidopsis* leaves lacking glucan water dikinase (*gwd*−/−), and liver glycogen to determine if this method can distinguish the difference between different phosphate levels. As predicted, the method detected the highest level of phosphate content in WT starch followed next by liver glycogen. The *gwd*−/− *Arabidopsis* starch contained no detectable G3P or G6P (**Fig. 5**). These data confirm the application of this novel GCMS-based method to quantify glucose-based polymers beyond mammalian glycogen and demonstrate that it can be used for the quantitation of plant starch.

**Fig. 5.**
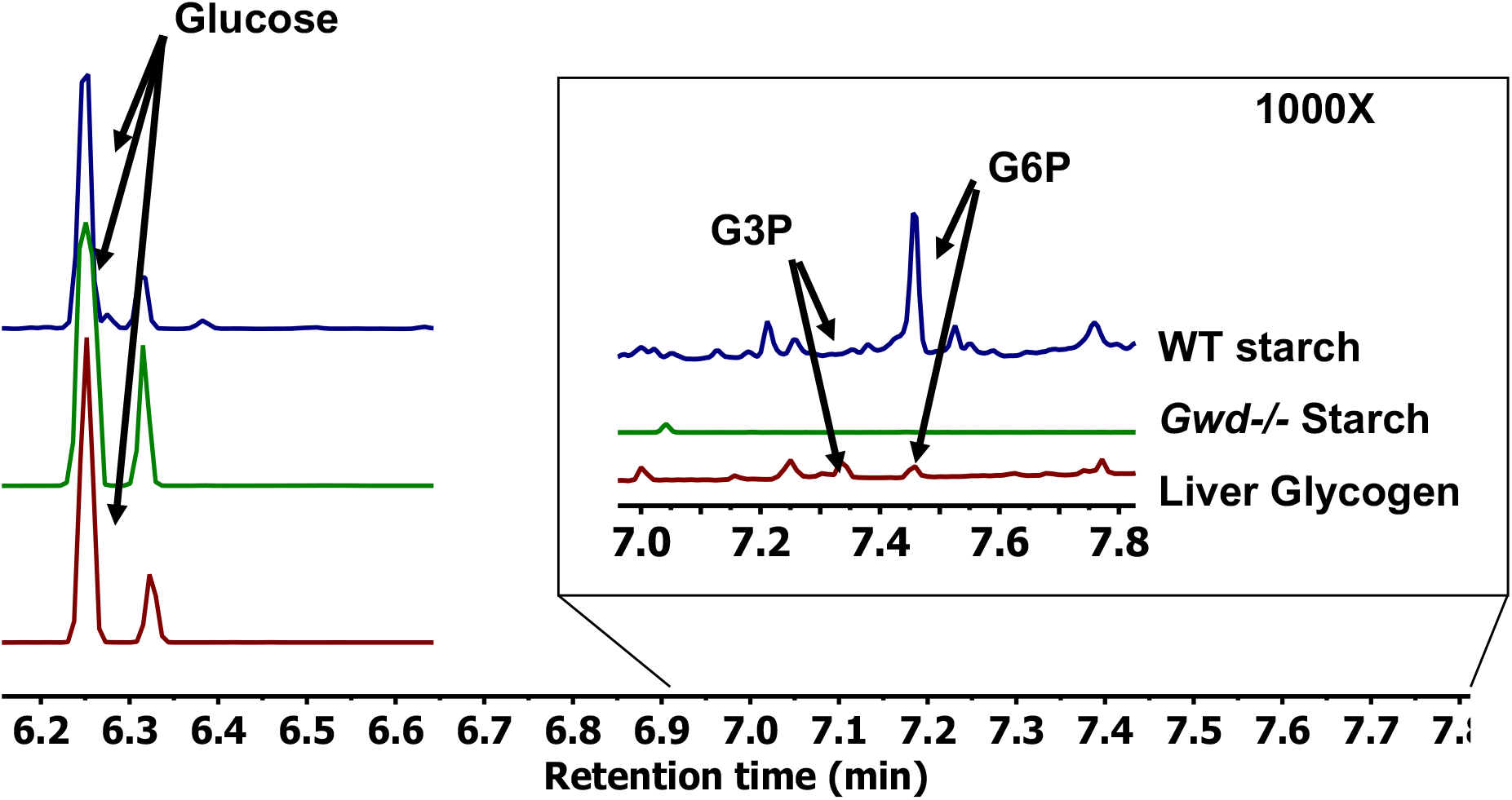
Comparison between plant starch and mammalian liver glycogen. Stacked spectra overlay between WT potato starch, *gwd*−/− *Arabidopsis* starch, and mammalian liver glycogen to demonstrate versatility of the approach. All three samples were standardized to total glucose. The region between 7-7.8 minutes demonstrates that WT plant starch contains 10-fold higher phosphate than liver glycogen, while no detectable phosphate was observed in the *gwd*−/− *Arabidopsis* starch.

## Discussion

GCMS instruments are routinely used in analytical chemistry and known for their resolution range, reproducibility, and durability. A new generation of autosampler-enabled GCMS units are designed to provide a workhorse-platform that offer unmatched consistency and performance for a wide range of analytical needs. Basic and clinical researchers have developed GCMS-based methods for profiling an impressive array of small metabolites to study perturbations in cellular metabolism (47,48) and to uncover disease biomarkers (49). In this study, we adapted the GCMS analytical platform for the analysis of glycogen architecture. We employed a two-step derivatization process utilizing MEOX and MSTFA yielded derivatives that were more phase volatile and chemically stable forms of glucose, G3P, and G6P (−6TMS;1MEOX). This procedure has been shown previously to dramatically reduce the boiling point, improve the thermal stability, and enhance the chromatography separation of metabolites. We demonstrate the distinct separation of glucose, G3P, and G6P using a 12-minute chromatography method with a 20 fmol to 20 nmol dynamic range. This method is extremely robust, medium-throughput (120 samples/day), and can be adapted for the characterization of plant starch and amylopectin.

Accurate quantitation of carbohydrate polymers and phosphate esters remains challenging due to their unique biochemical and biophysical properties. We developed a workflow to purify glycogen from only 20mg of mouse tissue and accurately quantified the glycogen-derived glucose and glucose phosphate esters using GCMS. Based on the dynamic range of GCMS, this method can be further adapted to utilize even less sample input. We demonstrate an unambiguous measurement of glycogen-derived glucose, G3P, and G6P from muscle and liver. Our results align with previously published results, but this new method utilizes far less tissue to purify the glycogen and allows simultaneous quantification of glucose, G3P, and G6P. The application of this method is further demonstrated by the analysis of wild type potato starch, liver glycogen, and *gwd*−/− starch (42). This analysis confirms previous results that *gwd*−/− starch lacks phosphate and demonstrates the range in defining G6P between plant starch and liver glycogen.

Glucose-2-phosphate (G2P) has been identified as a possible third monomeric phosphate ester in glycogen with a similar concentration to G3P and G6P by NMR (12). Interestingly, we observed an additional peak in liver glycogen with the identical fragmentation ion and close retention to G3P and G6P. Based on its retention time and m/z ratio, we hypothesize that this unknown peak is G2P. However, a G2P standard is not available for confirmation.

This method will allow the robust analysis of glycogen polymers that accumulate in human glycogen storage diseases (GSD). The GSDs are a unique collection of monogenic diseases that share the accumulation of aberrant glycogen aggregates. While each GSD yields a glycogen-like aggregate, the exact architecture of the aggregate is unknown for many of the GSDs. Understanding the glycogen architecture would assist in defining the mechanism of disease pathology and facilitate with assessment of future treatment efficacy. Similarly, starch phosphorylation impacts multiple aspects of starch industrial processing. Starch is both a first-generation biofuel and industrial feedstock for paper, textiles, adhesives, and plastics. Phosphorylation is the only known natural modification of starch, and it directly influences starch hydration, crystallinity, freeze-thaw stability, viscosity, and transparency, which are all central to industrial applications (50). This method would also allow rapid analysis of starch from multiple species and genotypes.

## Experimental Procedures

### Materials

Analytical standards of D-glucose from Sigma Aldrich (Cas# 50-99-7), Glucose-3-phosphate synthesized by Chiroblock, and D-Glucose-6-phosphate disodium salt, C6H11Na2O9P·xH2O, from Sigma Aldrich (CAS #3671-99-6) were used throughout the study. Wild-type potato starch from Sigma Aldrich (CAS #9005-25-8) was purchased to test versatility of the method. *gwd*−/− *Arabidopsis* starch was a generous gift from Drs. Sam Zeeman and Diana Santelia. Mice were housed in a climate-controlled environment with a 14/10-hour light/dark cycle (lights on at 0600 hours) with water and solid diet provided *ad libitum* throughout the study. The Institutional Animal Care and Use Committee at University of Kentucky has approved all of the animal procedures carried out in this study under PHS Assurance #A3336-01.

### Glycogen Purification

Mice were sacrificed by spinal dislocation, liver and muscle were removed immediately post-mortem, and washed once with PBS, twice with diH2O, blotted dry, and snap frozen in Liquid nitrogen. The frozen tissues were pulverized to 10 μm particles in liquid N2 using a Freezer/Mill Cryogenic Grinder (SPEX SamplePrep). Twenty milligrams of each pulverized tissue were extracted in 50% methanol/chloroform (V/V 1:1) and separated into polar (aqueous layer), lipid (chloroform layer) and protein/DNA/glycogen (interfacial layer) fractions. Glycogen is further purified from protein/DNA using 1.5ml of ice cold 10% trichloroacetic acid. The glycogen fraction and polar fraction were dried by vacuum centrifuge at 10^−3^ mBar for hydrolysis and derivatization.

### Glycogen and Starch Hydrolysis

Hydrolysis of glycogen and starch was performed by first resuspending the pellet in diH2O followed by the addition equal parts 2N HCl. Samples were vortexed thoroughly and incubated at 95 ^o^C for 2 hours. The reaction was quenched with 100% methanol with 40uM L-norvaline (as an internal control). The sample was then incubated on ice for at least 30 minutes. The supernatant was collected by centrifugation at 15,0000 rpm at 4°C for 10 minutes and subsequently dried by vacuum centrifuge at 10^−3^ mBar.

### Sample Derivatization

Dried hydrolyzed glycogen samples were derivatized by the addition of 20mg/ml methoxyamine in pyridine and sequential addition of N-methyl-trimethylsilyation (MSTFA). Both steps were incubated for 60 minutes at 60 °C with thorough mixing in between addition of solvents. The mixture was then transferred to a v-shaped glass chromatography vial and analyzed by the GCMS.

### GCMS Quantitation

Initial GCMS quantitation of the analytical standards, glycogen and starch are described previously from the Fiehn metabolomics GC method with the following modifications: the initial rate was held at 60 °C for 1 minute followed by 1-minute of run time, rising at 10 °C/minute to 325 °C and holding for 10 minutes followed by a run for 37.5 minutes (32,33). Adaptation of the Fiehn method is the following for rapid separation of Glucose-, G3P-, and G6P-6TMS;1MEOX: the initial rate was held at 60 °C for 1 minute followed by a 1-minute run. Ramp 1: rising at a rate of 60 °C/minute reaching 220 °C and running for 3.67 minutes. Ramp 2: rising at 30 °C/minute reaching 270 °C and running for 5.33 minutes. Ramp 3: rising at 30 °C/minute to 325 °C, holding for 5 minutes and running for 12.2 minutes followed by a post run of 60 °C for 1 minute. The electron ionization was set to 70eV. Select ion monitoring mode was used for quantitative measurement. Ions used for the metabolites that represent glycogen are: glucose (319, 364), G3P, and G6P (315, 357, 387), and L-norvaline (174). Batch data processing was performed Masshunter software from Agilent. Glucose, G3P, and G6P were standardized to procedural control, norvaline, before quantitated using standard curve generated from known standards.

### Statistics

Statistical analyses were carried out using GraphPad Prism. All numerical data are presented as mean ± SD except for xenograft tumor growth which is presented as mean ± S.E. Grouped analysis was performed using two-way ANOVA. Column analysis was performed using one-way ANOVA or t-test. A P-value less than 0.05 was considered statistically significant.

## Acknowledgements

We would like to thank Dr. Craig Vander Kooi and Gentry lab members for helpful discussion regarding the work.

